# Endogenous APOBEC3B Promotes CHK1 Inhibitor Sensitivity

**DOI:** 10.64898/2026.08.04.742831

**Authors:** Bojana Stefanovska, Benjamin C. Troness, Christopher D. Mullally, Bárbara de la Peña Avalos, Mahmoud A. Ibrahim, Yanjun Chen, Elisa Fanunza, Michael A. Carpenter, Reuben S. Harris

## Abstract

APOBEC3B (A3B) is a single-stranded DNA cytosine deaminase overexpressed in cancer, where it causes genomic DNA damage and mutations associated with tumor evolution. Enforced A3B overexpression triggers a dependency on the replication-stress response in different cellular models. However, whether endogenous A3B in cancer cells might yield a similar vulnerability is unclear. Here, we investigate how endogenous A3B expression and catalytic activity affect sensitivity to CHK1 inhibition, using two cancer cell lines, JHOC5 and U2OS. A3B-expressing cancer cells are sensitive to two chemically distinct CHK1 inhibitors, GDC-0575 and Prexasertib. CHK1 inhibitor sensitivity is reduced by *A3B* CRISPR knockout and restored by re-expressing wildtype A3B in knockout cells. Moreover, an endogenous A3B-E255A catalytic mutant generated by homology-directed repair phenocopies the reduced CHK1 inhibitor sensitivity of *A3B*-null cells, demonstrating a DNA deamination-dependent mechanism. CHK1 inhibition induces replication-associated DNA damage and cell-cycle perturbation dependent on A3B expression. As a result, A3B-expressing cells accumulate more pan-nuclear γH2AX, aberrant DNA-content profiles, and an expanded EdU-negative S-phase population, which are hallmarks of stalled DNA replication. In comparison, *A3B*-null and A3B-E255A cells retain defined cell-cycle distributions and are less sensitive to CHK1 inhibition. Together, these findings identify endogenous A3B-catalyzed deamination as a therapeutically actionable source of replication-associated DNA damage that renders tumor cells selectively dependent on CHK1 function.

**Statement of significance:** APOBEC3B causes mutations in cancer cells and simultaneously imposes DNA replication stress. This combines to sensitize tumor cells to chemical inhibitors of the DNA damage response kinase CHK1.

## Introduction

The apolipoprotein B mRNA-editing enzyme catalytic polypeptide-like 3 (APOBEC3) family of cytidine deaminases was initially discovered as part of the innate antiviral immune response, restricting the replication of retroviruses, retrotransposons, and other viruses with susceptible single-stranded DNA (1–3). Unexpectedly, the same family of enzymes has become one of the most prevalent endogenous sources of somatic mutation in human cancer, with APOBEC3 mutational signatures detected in approximately 70% of cancer types (4). APOBEC3 mutagenesis is characterized by two canonical single base substitution (SBS) mutation signatures, SBS2 and SBS13, causing C-to-T and C-to-G substitutions, respectively, in 5′-TCW motifs (5–7). APOBEC activity also generates localized clusters of mutations known as *didyma*, *omikli*, and *kataegis*, which are respectively comprised of clusters of 2, 2-3, and ≥4 strand-coordinated APOBEC signature mutations and associated respectively with nucleotide excision repair, mismatch repair, and recombination repair/R-loop formation (8–11). APOBEC3 mutagenesis is also DNA replication-associated and able to access both leading- and lagging-strand intermediates (with bias toward the latter substrates) (12–14). Through these mechanisms, APOBEC3 mutagenesis promotes tumor evolution including the acquisition of mutations that confer therapy resistance and drive metastases (15–21).

APOBEC3A (A3A) and APOBEC3B (A3B) are the prime contributors to the overall APOBEC mutational signature in human cancer (5–7,22–25). Moreover, A3B is the only enzyme that is constitutively nuclear (26–29). A3B deaminates cytosine to uracil in single-stranded DNA (ssDNA), and the resulting lesions can be replicated directly or processed by DNA repair pathways into base substitution mutations (SBS2 and SBS13), DNA strand breaks, and chromosomal instability. A3B is upregulated and/or aberrantly expressed in several tumor types, and its experimental expression in mice recapitulates APOBEC signature mutations as well as *didyma* and *kataegis* (5,8,21,22). Although A3A also contributes to the overall APOBEC mutation signature in cancer, the sustained high levels of expression, nuclear localization, and tumor-promoting activity of A3B make it a potentially exploitable therapeutic target.

A3B activity is closely linked to DNA replication stress (30–37). Stalled or perturbed replication forks expose stretches of ssDNA that serve as substrates for deamination by APOBEC3 enzymes. Thus, A3B-mediated deamination and processing of the resulting uracil lesions can impede fork progression, generate abasic sites and single- and double-stranded DNA breaks, and increase the amount of replication-associated ssDNA. Abasic lesions at replication forks may also promote RPA exhaustion and replication stress and, together with ATR inhibition, can trigger cell death (34,36). A3B can also bind, deaminate, and regulate R-loops, DNA/RNA hybrid structures that form by transcripts re-annealing to genic DNA template strands (10,38). R-loops can also interfere with DNA replication when they persist or resolve aberrantly (39). In melanoma models, A3B depletion reduces R-loop abundance and replication-stress signaling, whereas A3B overexpression increases both phenotypes (33). Expression of Ribonuclease H1, which resolves R-loops, substantially reverses these effects (33). More recent work has further shown that, following ATR inhibition, A3B-mediated uracil formation can promote APE1-dependent DNA cleavage, PARP1 trapping, and replication-fork breakage (36). Taken together, these observations suggest that in addition to signature single base substitution mutations, A3B may also contribute to real-time replication stress in cells where it is overexpressed.

In this study, we generate isogenic cancer cell models with null and high endogenous A3B expression levels to test whether endogenous levels of this enzyme also confer sensitivity to inhibition of replication-stress responses. We show that A3B-proficient cells exhibit preferential sensitivity to CHK1 inhibition, whereas inhibition of ATR or WEE1 produces weaker and more context-dependent effects. Sensitivity to CHK1 inhibition also requires A3B catalytic activity. Moreover, in presence of CHK1 inhibition, A3B activity causes a disruption in cell cycle progression and a pan-nuclear accumulation of γH2AX. These results combine to indicate that endogenous A3B activity creates a functional dependence on CHK1-mediated replication stress responses.

## Materials and methods

### Cell Culture

U2OS cells were purchased from ATCC (HTB-96), 293T cells were purchased from ATCC (CRL-3216), and JHOC5 cells were obtained from Tian-Li Wang at Johns Hopkins University. Cells were maintained in RPMI 1640 medium (Gibco, 11875093) supplemented with 10% fetal bovine serum (BioWest, 058N24) and cultured at 37°C in a humidified incubator with 5% CO_2_. All cell lines were confirmed to be Mycoplasma-free and were tested quarterly using the MycoAlert Mycoplasma Detection Kit (Lonza, LT07).

### Generation of *A3B*-knockout and *A3B-E255A* knock-in models

The U2OS *A3B* knockout clone was generated as described (10). The catalytically inactive A3B E255A mutant was generated by CRISPR/Cas9-mediated homology-directed repair. Glutamate 255 was converted to alanine using an sgRNA targeting exon 6 of *A3B* (5′-AGAAGCGCAGCTCCGCATGG-3′) together with a single-stranded oligodeoxynucleotide (ssODN) donor carrying the corresponding GAG-to-GCG codon substitution, flanked by 42-bp (5′) and 40-bp (3′) homology arms.

Ribonucleoprotein (RNP) complexes were assembled by combining Cas9 protein with the A3B-targeting sgRNA prior to electroporation and were delivered using the Neon Transfection System (Thermo Fisher Scientific, NEON18). For each reaction, 2.0× 10^5^ cells were resuspended in 10 µL of Neon Resuspension Buffer R and mixed with 1,250 ng Cas9 protein, 7.5 pmol sgRNA, and 100 pmol of the E255A ssODN donor. Cells were electroporated at 1,400 V with a 15-ms pulse width for four pulses, then immediately transferred to a 24-well plate containing prewarmed, antibiotic-free complete RPMI medium and cultured at 37°C in a humidified atmosphere containing 5% CO_2_. Forty-eight hours after electroporation, cells were plated by limiting dilution into 96-well plates to isolate single-cell clones. Clones were screened by immunoblotting and an oligo-based C-to-U deamination assay. Those retaining A3B protein expression but lacking detectable catalytic activity were kept and further validated by PCR amplification across the sgRNA target region followed by Sanger sequencing to confirm the E255A substitution.

For JHOC5, the parental cell line was first single-cell cloned to establish a mother line, which was then transduced with lentivirus produced from pLentiCRISPR encoding an sgRNA targeting either *A3B* or *lacZ* (control) (40). Transduced cells were selected with 1.5 µg/ml puromycin for 72 h and then seeded for single-cell cloning. Resulting clones were screened for *A3B* knockout by immunoblotting and validated as above. After validation, candidate clones were transfected with Cre-recombinase plasmid, to remove Cas9 and puromycin resistance, as described (40). All sgRNAs, donor sequences, and validation primers are listed in **Supplementary Table 1**.

### Generation of A3B overexpression models

JHOC5 and U2OS wildtype and *A3B* knockout cells were engineered to stably express A3B using collapsing retroviral (CRV 1.0) vectors, as described (41). Retroviral particles were produced by transient transfection of 293T cells. Briefly, 293T cells were transfected with the CRV-A3×3B retroviral plasmid together with the appropriate packaging and envelope plasmids using LT-1 transfection reagent. Viral supernatants were harvested 72 h after transfection and purified through a 0.45-µm filter. Immediately after filtration, target cells were transduced in the presence of 4 µg/mL polybrene. Forty-eight hours after transduction, cells were selected with 1.5 µg/mL puromycin (Gibco, A1113803) to establish stable bulk populations. Following selection, A3B expression was assessed by immunoblotting and catalytic activity was confirmed using an oligo cleavage deaminase activity assay, as described below.

### Drug treatments

The following small-molecule inhibitors were used: ATR inhibitor AZD6738 (Selleck Chemicals, S7693), WEE1 inhibitor ZNL-02-096 (Tocris, 7240), and CHK1 inhibitors GDC-0575 (Selleck Chemicals, S8526) and Prexasertib (Selleck Chemicals, S7178). All inhibitors were purchased as powders and dissolved in 100% DMSO to generate stock solutions.

### Immunoblotting

JHOC5 and U2OS cells were seeded at 9375 cells/cm^2^ in 6-well plates. 24 h after seeding, the growth medium was replaced with 2 mL fresh medium containing DNA damage response inhibitor or equivalent volume of DMSO as a vehicle control. Cells were incubated 72 h with the respective inhibitor. After treatment, cells were flash-frozen by briefly submerging the bottom of the plate in a dry ice–ethanol bath and lysed in Pierce™ RIPA buffer (ThermoFisher Scientific, 89901) supplemented with 1× cOmplete™ EDTA-free protease inhibitor cocktail (Roche, 04906837001) and PhosSTOP phosphatase inhibitor cocktail (Roche, 05056489001). Total protein was quantified using a bicinchoninic acid (BCA) protein assay kit (EMD Millipore, 71285-3). Equal amounts of protein were resuspended in SDS-PAGE loading buffer (125 mM Tris pH 6.8, 5% SDS, 30% glycerol, 100 mM dithiothreitol [DTT], and Orange G dye) and denatured at 98 °C for 10 min.

Proteins were resolved by SDS-PAGE on a 4–20% gel and transferred to an Immobilon-FL polyvinylidene difluoride membrane (Millipore, IPFL00010) using a Bio-Rad Trans-Blot Turbo apparatus according to the manufacturer’s protocol. Transfer efficiency was confirmed by Ponceau staining (Sigma-Aldrich, P7170). Membranes were blocked in 1× casein buffer for 1 hour at room temperature. Primary and secondary antibodies are listed in **Supplementary Table 1**. Primary antibodies were diluted in EveryBlot Blocking Buffer (Bio-Rad, 12010020) for commercial antibodies or in 1× casein buffer for the anti-A3A/B/G antibody, and membranes were incubated overnight at 4 °C with gentle rocking. Membranes were then washed three times in PBS containing 0.1% Tween-20 (PBS-T) and once in PBS, 5 min each. Secondary antibodies were diluted in 1× casein buffer and incubated with the membranes for 1 hour at room temperature with gentle rocking. HRP-conjugated secondary antibodies were developed using SuperSignal™ West Femto Maximum Sensitivity Substrate (Thermo Scientific, 34096) and fluorescent secondary antibodies were imaged on a LI-COR Odyssey-M system.

### Oligonucleotide-based DNA C-to-U activity assays

Single-stranded DNA cytosine deaminase activity in whole-cell lysates was measured using a fluorescence-based oligonucleotide cleavage assay (5,10). JHOC5 and U2OS cells were lysed in HED buffer containing 25 mM HEPES, 5 mM EDTA, 10% glycerol, 1 mM DTT, and 1× cOmplete protease inhibitor cocktail. Cleared whole-cell lysate equivalent to 7.5 µg total protein was used for each reaction. Each 20 µL reaction contained 4 pmol of a 3′ FAM-labelled single-stranded DNA substrate (5′-ATTATTATTATTCTAATGGATTTATTTATTTATTTATTTATTT-FAM), 0.025 U uracil-DNA glycosylase (UDG; NEB, M0280), and 1.75 U RNase A (NEB, M0314). Reactions were incubated 2 h at 37°C. Following incubation, samples were treated with 100 mM NaOH and heated 10 min at 95°C to cleave the DNA backbone at abasic sites generated following UDG-mediated removal of uracil. An equal volume of 2X formamide sample buffer (80% formamide, 0.05% bromophenol blue, and 0.01% xylene cyanol in TBE) was then added to each reaction and incubated another 5 minutes at 98°C. Substrate and cleavage products were separated on 15% denaturing TBE–urea polyacrylamide gels, run at 12 Watts for 45 min, and FAM fluorescence was imaged using a LI-COR Odyssey M imaging system.

### Cell viability measurements

JHOC5 and U2OS cells were seeded at 9375 cells/cm^2^ in 96-well plates. 24 h after seeding, the media was removed and replaced with 100 μL of fresh growth media containing DNA damage response inhibitors or equivalent volume of DMSO as a vehicle control. Cells were cultured in the presence of inhibitors for 72 h, after which viability was assessed using the CellTiter-Glo luminescent cell viability assay (Promega, G8461). Briefly, drug-containing medium was removed and replaced with 50 µL PBS and 50 µL CellTiter-Glo reagent per well. Plates were orbitally shaken for 2 min and then incubated for 15 min on a rocker to allow cell lysis and signal stabilization. A 50 µL aliquot from each well was transferred to a white-walled 96-well plate, and luminescence was measured using a Tecan Spark multimode microplate reader. At least three biological replicates were performed for each condition. Data analysis was performed with GraphPad Prism (v11.0.0).

### Colony formation assays

JHOC5 and U2OS cells were seeded at 400 and 800 cells per well in 6-well plates with 2 mL of complete growth medium in each well. After 24 h, the medium was removed and replaced with 2 mL of fresh growth medium containing the indicated DNA damage response inhibitor or an equivalent volume of DMSO as a vehicle control. Inhibitor-containing medium was replaced every 4 days. JHOC5 cells were cultured for 8 days and U2OS cells for 10 days after treatment to allow formation of countable colonies. Cells were then washed with PBS, fixed with 80% ethanol, and stained with crystal violet staining solution containing 50% methanol and 0.5% w/v crystal violet (Fisher Chemical, C581). Wells were washed twice with water and allowed to dry before imaging using a LI-COR Odyssey M imaging system. Colonies containing 50 or more cells were counted manually. At least three biological replicates were performed for each condition. Data were analyzed using GraphPad Prism v11.0.0.

### Cell cycle analyses

Cell cycle distribution was assessed by propidium iodide (PI) staining alone or by combined 5-ethynyl-2′-deoxyuridine (EdU, Invitrogen, C10635) incorporation and PI staining. For PI-only cell cycle analysis, cells were seeded at 9375 cells/cm^2^ in 6-well plates. After 24 h, the medium was removed and replaced with fresh growth medium containing the indicated DNA damage response inhibitor or an equivalent volume of DMSO as a vehicle control. At 24 h post-treatment, cells were harvested, washed with PBS, and fixed in 250 µL ice-cold 100% methanol for 30 min on ice. Fixed cells were then washed three times with PBS and resuspended in 400 µL DNA staining solution containing 20 µg/mL PI, 200 µg/mL RNase A, and 0.1% Triton X-100 in PBS. Cells were incubated for 30 min at room temperature in the dark and analyzed using a LSRFortessa X-20 flow cytometer equipped with a high-throughput sampler (BD Biosciences, 656385).

For combined EdU and PI staining, cells were seeded and treated as described above. At 24 h post-treatment, EdU was added directly to the culture medium at a final concentration of 10 µM, and cells were incubated for 1 h before harvesting. Cells were then collected, fixed in 250 µL ice-cold 100% methanol for 30 min on ice, and washed three times with PBS. EdU and its incorporation was assessed using the Click-iT Plus EdU Alexa Fluor 647 Flow Cytometry Assay Kit (Invitrogen, C10635). Fixed cells were permeabilized in 1×Click-iT saponin-based permeabilization and wash reagent for 15 min, then incubated with the Click-iT Plus reaction cocktail for 30 min at room temperature protected from light. Cells were washed again with permeabilization and wash reagent and resuspended in 400 µL DNA staining solution containing PI, RNase A, and Triton X-100. After incubation for 30 min at room temperature in the dark, samples were analyzed using a BD LSRFortessa X-20 flow cytometer equipped with a high-throughput sampler. Flow cytometry data were analyzed using FlowJo v10 (BD Biosciences).

### Immunofluorescence microscopy assays

JHOC5 and U2OS cells were seeded at 9375 cells/cm^2^ in 4-chamber tissue-culture slides containing 1 mL of complete growth medium. After 24 h, the medium was removed and replaced with 1 mL of fresh growth medium containing the indicated DNA damage response inhibitor or an equivalent volume of DMSO as a vehicle control. At 24 h post-treatment, medium was removed and cells were washed with PBS before fixation with 4% paraformaldehyde for 10 min. Cells were washed twice with PBS and permeabilized with 0.2% Triton X-100 (Thermo Scientific, A16046) in PBS for 10 min. Cells were then blocked for 10 min in blocking buffer containing 5% goat serum (Gibco, 16210064) and 0.1% Triton X-100 in PBS. Cells were incubated overnight at 4°C on a rocker with mouse anti-γH2AX S139 primary antibody diluted 1:500 in blocking buffer (Millipore Sigma, 05-636). Cells were then incubated for 1 h at room temperature with goat anti-mouse Alexa Fluor 647 secondary antibody diluted 1:1000 in blocking buffer (Invitrogen, A32728). Following staining, cells were washed with PBS and mounted with coverslips using Ibidi Mounting Medium with DAPI (Ibidi, 50011). Images were acquired using a 60× objective on a Nikon Eclipse Ti2 microscope.

Image segmentation to identify cell nuclei and nuclei with pan-nuclear γH2AX was performed using CellProfiler version 4, an open-source image analysis program (42). To identify nuclei, the “RescaleIntensity” function was used to enhance visibility, and the “IdentifyPrimaryObjects” function was used on images in the DAPI channel. To identify nuclei with pan-nuclear γH2AX expression, the “RescaleIntensity” function was used to enhance visibility and “IdentifyPrimaryObjects” was used on images in the Cy5 channel. The identified objects were then related back to the identified nuclei with the “RelateObjects” function. Data were analyzed using GraphPad Prism v11.0.0.

### Statistical analyses

Data analyses were performed using GraphPad Prism. All data represent results from 3 independent experiments and are presented as mean ± SD. Statistical significance was defined as p < 0.05. Statistical tests used and corresponding p-values for each experiment are provided in the relevant figure legend.

## Results

### Endogenous A3B confers a selective vulnerability to CHK1 inhibition

The DNA cytosine deaminase A3B is overexpressed in many different tumor types and cancer cell lines, and rarely in normal tissues and normal-like cell lines (22–24,43–45). A3B deaminates genomic cytosines to uracils, resulting in abasic sites and DNA breaks, and its enforced overexpression can trigger chronic replication stress and replication-associated DNA damage (32,46). To further investigate the link between A3B activity and replication stress in cancer cells, CRISPR was used to generate *A3B*-null derivatives of two A3B-high cell lines, the ovarian cancer cell line JHOC5 and the osteosarcoma cell line U2OS. These two lines were chosen for mechanistic studies here due to pathologically high (tumor-like) endogenous A3B protein levels and amenability to immunofluorescence microscopy and molecular assays. Candidate A3B-knockout clones were identified by loss of A3B protein on immunoblotting and validated by Sanger sequencing across the gRNA target site to confirm frameshifting indels. DNA sequencing confirmed frameshift mutations in *A3B* exon 3, including a +1 T insertion in JHOC5 and +1 T or +1 A insertions in U2OS (**Fig. 1A**). Immunoblotting confirmed total loss of A3B protein expression and DNA deaminase activity assays showed a corresponding loss of activity in whole cell extracts (**Fig. 1B**).

**Figure 1.**
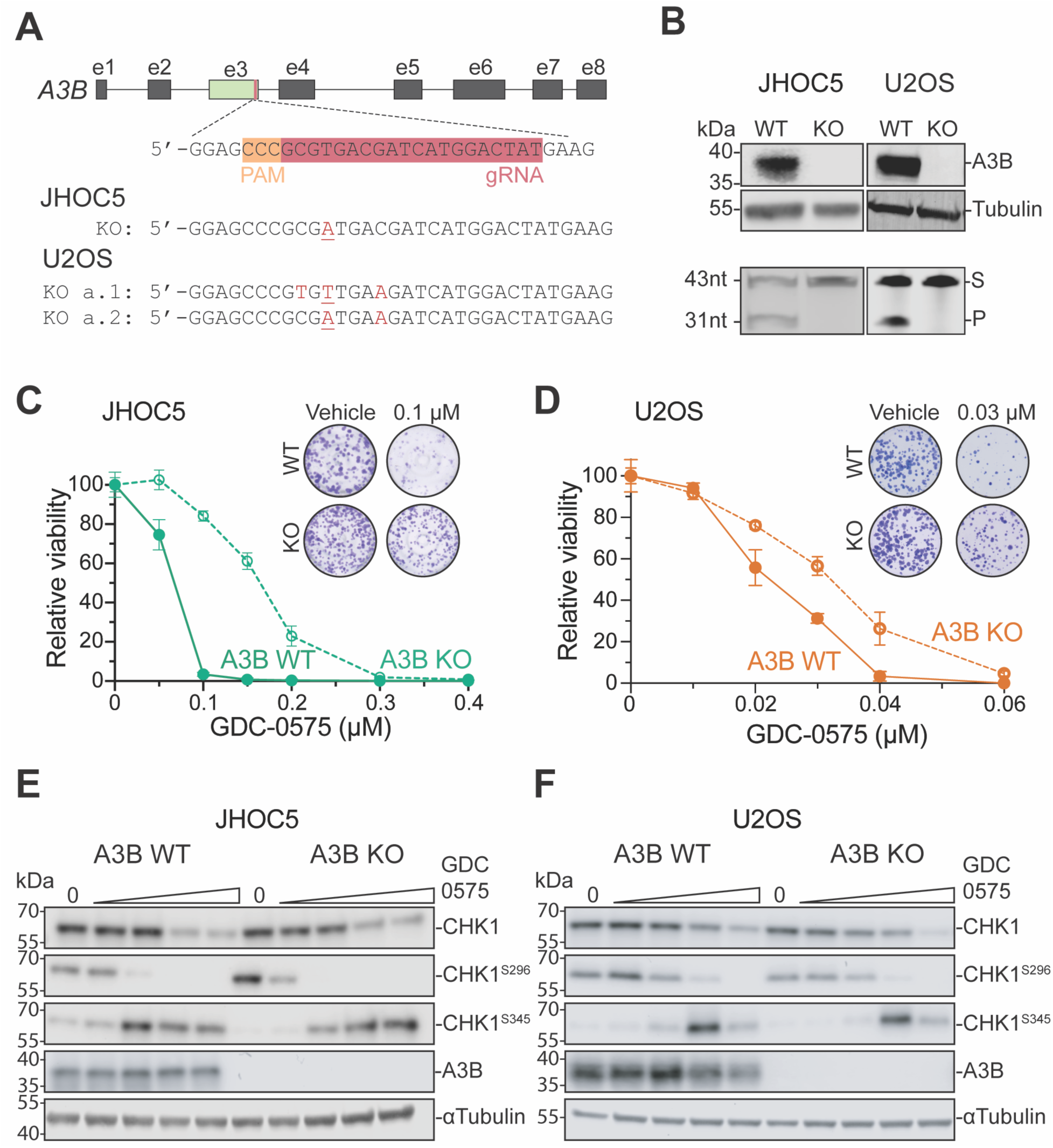
APOBEC3B-proficient cells are sensitive to the CHK1 inhibitor GDC-0575. **A**, Schematic of the *A3B* knockout strategy. *A3B* exon 3 was targeted for disruption by CRISPR as indicated, and the resulting mutant genotypes are shown below for each cell line (mutations are red and underlined). **B**, Immunoblots of A3B expression in WT and *A3B* KO JHOC5 and U2OS cells (top panel). DNA deaminase activity of whole-cell lysates from the indicated cell lines (S, substrate; P, product; bottom panel). **C** and **D**, Dose-response analysis of *A3B* WT versus KO JHOC5 and U2OS cells, respectively, following treatment with the indicated concentrations of the CHK1 inhibitor GDC-0575. Relative viability was measured after drug treatment and normalized to vehicle-treated controls. Representative colony-formation images are shown. Data are mean ± SD, n = 3 independent experiments. (P-value <0.0001 for WT vs KO by extra sum-of-squares F-test). **E**, Immunoblot analyses of CHK1 signaling in *A3B* WT and KO JHOC5 following treatment with 0, 0.005, 0.05, 0.5, and 5 µM GDC-0575. **F**, Immunoblot analyses of CHK1 signaling in *A3B* WT and KO U2OS following treatment with 0, 0.001, 0.01, 0.1, and 1 µM GDC-0575.

As CHK1 is the principal effector kinase downstream of ATR, two chemically distinct CHK1 inhibitors, GDC-0575 and Prexasertib, were used to probe for a possible association with A3B and replication stress. Both inhibitors bind selectively to the ATP pocket of CHK1, with Prexasertib also capable of binding to CHK2 at higher concentrations (47,48). Interestingly, both A3B-expressing parental cell lines exhibited a greater sensitivity to GDC-0575 than their otherwise isogenic *A3B*-null counterparts (**Fig. 1C** and **D**). This A3B-dependent sensitivity is clear in cell viability assays such as CellTiter-Glo (**Supplementary Fig. 1A** and **B**), but it is even more pronounced in gold-standard colony formation assays (dose response curves and representative images of stained colonies in insets of **Fig. 1C** and **D**). Similar results were obtained with Prexasertib, though phenotypes in the A3B-expressing JHOC5 cells were much stronger than those for U2OS cells (**Supplementary Fig. 1C** and **D**; **Discussion**).

To confirm the pharmacologic activity of both inhibitors, immunoblotting for total CHK1 and established markers of CHK1 pathway activity was performed following treatment with increasing doses of GDC-0575 or Prexasertib. As expected, in both JHOC5 and U2OS cells, GDC-0575 and Prexasertib treatment reduced CHK1 S296 autophosphorylation, consistent with on-target inhibition of CHK1 kinase activity (**Fig. 1E** and **F**; **Supplementary Fig. 1E** and **F**). CHK1 S345 phosphorylation, an ATR-dependent marker of upstream checkpoint activation, was also assessed and CHK1 inhibition led to increased S345 phosphorylation in both cell lines, which is consistent with an accumulation of replication stress signaling upon CHK1 blockade. Interestingly, in JHOC5 cells, basal CHK1 S345 phosphorylation appeared lower in *A3B*-null cells than in the A3B-expressing parental cells, although this difference was not observed in the U2OS cells (**Fig. 1E** and **Supplementary Fig. 1E**).

### A3B catalytic activity is essential for sensitivity to CHK1 inhibitors

A3B has both catalytic and non-catalytic activities, with single-stranded DNA C-to-U deamination being its canonical activity attributable to the catalytic C-terminal domain and single-stranded DNA and RNA binding activity being attributable to both the non-catalytic N-terminal domain as well as the C-terminal catalytic domain (49,50). Nucleic acid binding is a pre-requisite for deamination, but it is also easy to envisage models where the strong binding activity of A3B may alone be sufficient to impose replication stress. To investigate whether CHK1 inhibitor sensitivity is driven by a deamination-dependent or -independent mechanism, CRISPR-HDR was used to generate an isogenic U2OS knock-in cell line carrying the catalytically inactive A3B E255A mutation (**Fig. 2A**). Successful generation of the mutant line was confirmed by Sanger sequencing, immunoblotting, and A3B deamination activity assays (**Fig. 2B**).

**Figure 2.**
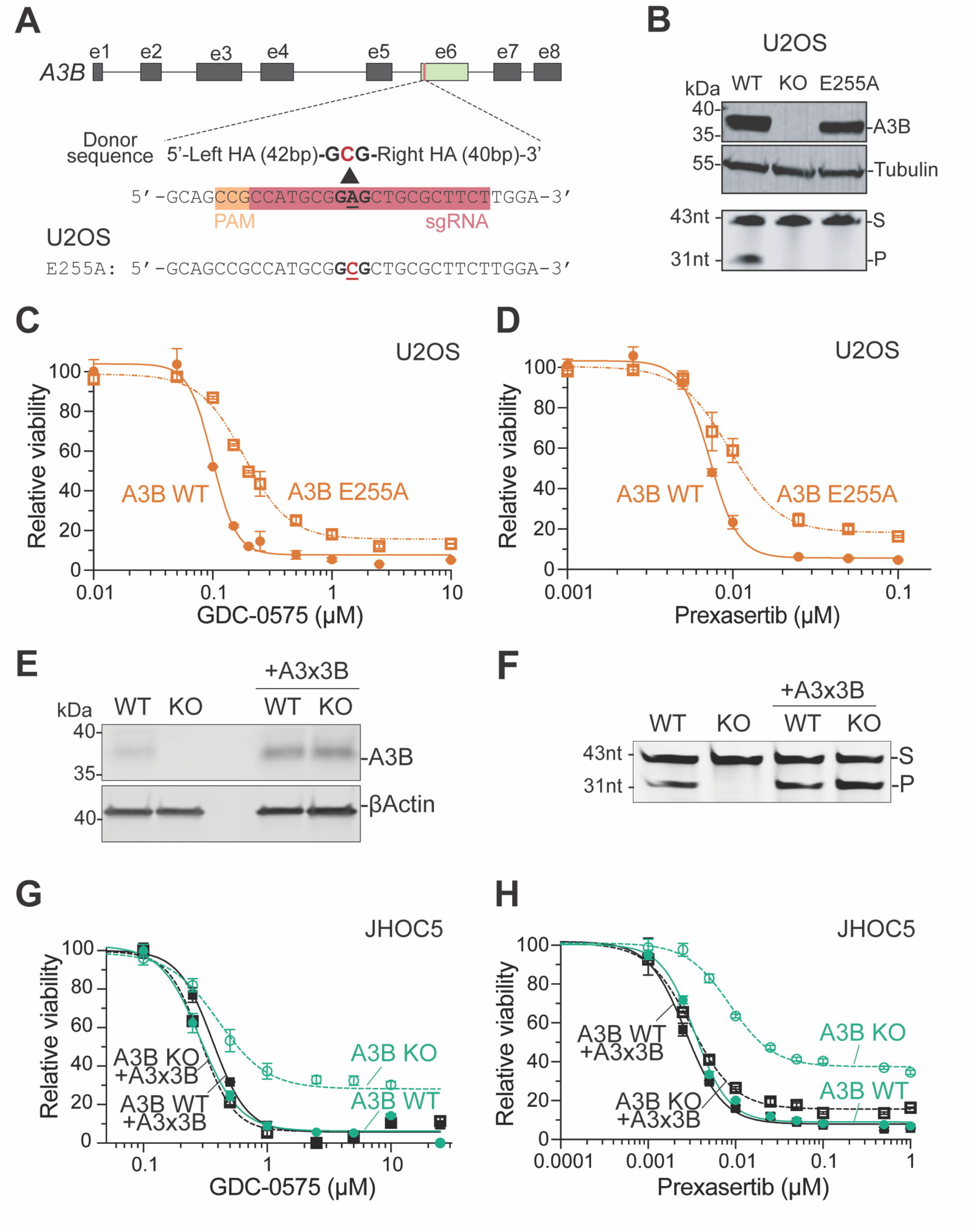
APOBEC3B catalytic activity promotes sensitivity to CHK1 inhibition. **A**, Schematic of the CRISPR-mediated homology-directed repair strategy to change Glu255 (GAG) to Ala (GCG) in *A3B* exon 6 in U2OS cells (mutation is red and underlined). **B**, Anti-A3B immunoblot showing protein expression in WT and E255A but not in KO U2OS cells (top panel). DNA deaminase activity of whole-cell lysates from the indicated cell lines (S, substrate; P, product; bottom panel). **C** and **D**, Dose-response analysis of *A3B* WT versus A3B-E255A expressing U2OS cells following treatment with GDC-0575 and Prexasertib, respectively. Relative viability was quantified using CellTiter-Glo 72 h after drug treatment (normalized to vehicle-treated controls; mean ± SD of 3 independent experiments; P-value <0.0001 by extra sum-of-squares F-test). **E**, Anti-A3B immunoblot demonstrating CRV-mediated expression of exogenous A3B in WT and KO JHOC5 cells **F**, DNA C-to-U deamination activity assay demonstrating CRV-mediated expression of exogenous A3B in WT and KO JHOC5 cells (S, substrate; P, product). **G** and **H**, Dose-response analysis of untransduced or CRV-transduced *A3B* WT versus *A3B* KO JHOC5 cells following treatment with GDC-0575 and Prexasertib, respectively. Relative viability was quantified using CellTiter-Glo 48 h after drug treatment (normalized to vehicle-treated controls; mean ± SD of 3 independent experiments; P-value <0.0001 for WT vs KO by extra sum-of-squares F test; all other comparisons are not significant).

Compared with A3B WT U2OS cells, A3B E255A knock-in cells exhibited reduced sensitivity to both GDC-0575 and Prexasertib measured by CellTiter-Glo (**Fig. 2C-D)**. The reduced sensitivity of the E255A knock-in cells was similar to that observed in A3B KO cells, indicating that catalytic inactivation of A3B phenocopies the *A3B*-null allele in this context.

To further investigate if A3B is necessary for CHK1 inhibitor sensitivity, the *A3B*-null JHOC5 and U2OS clones were engineered by collapsing retrovirus (CRV) transduction to stably express wildtype A3B (**Methods**). Ectopic expression of A3B was confirmed by immunoblotting, and deaminase activity assays showed that the reintroduced protein was catalytically active (**Fig. 2E-F**, **Supplementary Fig. 2A-B**). Viability measurements by CellTiter-Glo confirmed again that *A3B*-null cells are less sensitive to CHK1 inhibition. In contrast, the reintroduction of A3B re-sensitized cells to CHK1 inhibition in a manner comparable to the parental cell line with WT A3B. This effect was more pronounced in the JHOC5 cell line (**Fig. 2G-H**), in comparison to the U2OS cell line (**Supplementary Fig. 2C-D**). Altogether, these results indicated that A3B-catalyzed DNA deamination promotes CHK1 inhibitor sensitivity.

### Endogenous A3B confers moderate sensitivity to ATR and WEE1 inhibitors

To test whether the A3B-associated CHK1 vulnerability described above is specific to CHK1, inhibition of the upstream kinase ATR and the downstream G2/M checkpoint kinase WEE1 was assessed for comparison. ATR is responsible for activating CHK1 in response to replication stress, whereas WEE1 restrains CDK1 activity and prevents premature entry into mitosis (51,52). Here, the viability of isogenic A3B-expressing and *A3B*-null JHOC5 and U2OS cells was quantified following treatment with the ATR inhibitor AZD6738 or the WEE1 degrader ZNL-02-096. In contrast to the robust sensitivity of A3B-expressing cells to CHK1 inhibitors, ATR and WEE1 inhibition produced weaker effects (**Supplementary Fig. 3A-F**).

Treatment with AZD6738 for 72 h reduced viability measured with CellTiter-Glo in both JHOC5 and U2OS cells, confirming drug activity, but it did not trigger higher sensitivity in A3B-expressing cells compared to *A3B*-null counterparts (**Supplementary Fig. 3A**). Similar results were observed also in colony formation assays. Prolonged AZD6738 exposure produced decrease in colony formation capacity in both cell lines, with a modest increase in the sensitivity of A3B expressing cells compared to *A3B*-null cells (**Supplementary Fig. 3C-D**). These results suggested that ATR inhibition may not phenocopy the stronger A3B-dependent sensitivity to CHK1 inhibition described above.

Treatment with the WEE1 degrader ZNL-02-096 for 72 h caused a modest but significant A3B-dependent decrease in viability in JHOC5 cells and, in contrast, no clear A3B-dependent difference in U2OS cells (**Supplementary Fig. 3B**). In colony formation assays, prolonged ZNL-02-096 exposure produced modest but significant differences between A3B-expressing and *A3B*-null cells in both models, though these effects were again less pronounced than with CHK1 inhibitors (**Supplementary Fig. 3E-F**). Therefore, in these two models, A3B expression and activity are more consistently associated with responses to CHK1 inhibition than with responses to ATR or WEE1 inhibition.

### A3B-expressing cells accumulate pan-nuclear γH2AX following CHK1 inhibition

JHOC5 and U2OS cells were stained for γH2AX to investigate whether the increased sensitivity to CHK1 inhibition is associated with DNA damage. Pan-nuclear γH2AX is a marker of widespread replication-associated DNA damage (53). After 24 h of treatment with the CHK1 inhibitor GDC-0575, A3B-expressing JHOC5 cells showed a marked increase in pan-nuclear γH2AX staining at both IC_50_ and IC_95_ concentrations (**Fig. 3A**). In contrast, *A3B*-null cells showed substantially lower levels of pan-nuclear γH2AX under the same treatment conditions. Re-expression of A3B in *A3B*-null cells through retroviral transduction restored elevated pan-nuclear γH2AX at these two GDC-0575 concentrations, supporting a role for A3B in promoting CHK1 inhibitor-associated DNA damage. Similar results were observed in JHOC5 cells treated with the CHK1 inhibitor Prexasertib (**Fig. 3B**).

**Figure 3.**
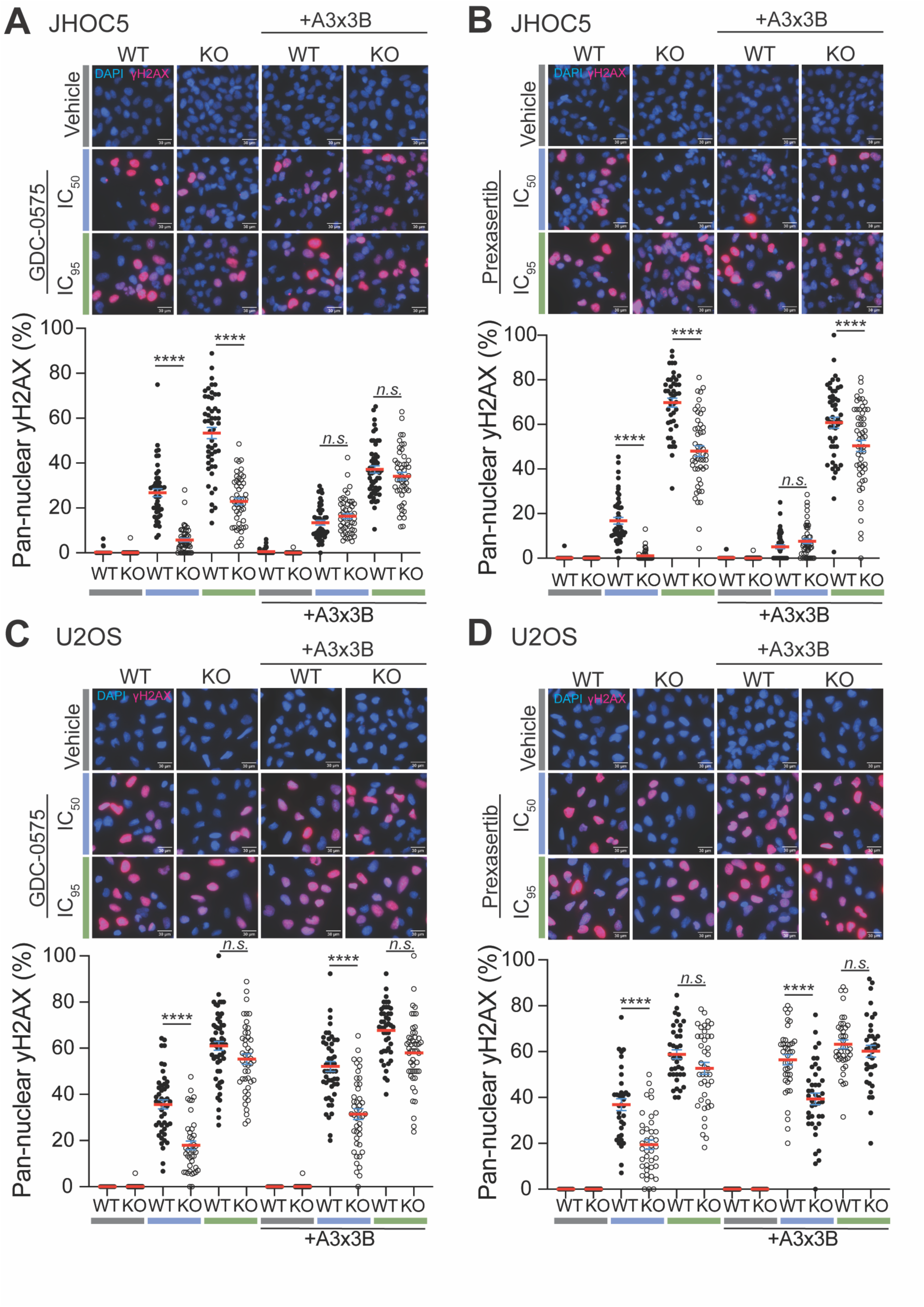
A3B expression promotes pan-nuclear γH2AX accumulation following CHK1 inhibition. **A** and **B**, Representative images of JHOC5 *A3B* WT and *A3B* KO cells treated with GDC-0575 at IC_50_ (0.15 µM) or IC_95_ (0.25 µM) concentrations, or Prexasertib at IC_50_ (0.005 µM) or IC_95_ (0.025 µM) concentrations. (30 μm scale). For quantification, each data point represents the percentage of cells with pan-nuclear γH2AX within single microscopy image (field); horizontal bars indicate the mean ± SEM of >40 fields per condition; significance assessed by two-tailed unpaired Student’s t-test (****, P < 0.0001; *n.s*., not significant). **C** and **D**, Representative images of U2OS *A3B* WT and *A3B* KO cells treated with GDC-0575 at IC_50_ (0.085 µM) or IC_95_ (0.25 µM) concentrations, or Prexasertib at IC_50_ (0.003 µM) or IC_95_ (0.0065 µM) concentrations (30 μm scale). For quantification, each data point represents the percentage of cells with pan-nuclear γH2AX within single microscopy image (field); horizontal bars indicate the mean ± SEM of >40 fields per condition; significance assessed by two-tailed unpaired Student’s t-test (****, P < 0.0001; *n.s*., not significant).

A3B-expressing U2OS cells treated with GDC-0575 showed increased pan-nuclear γH2AX compared with *A3B*-null cells at the IC_50_ concentration, whereas no significant difference was observed at the IC_95_ concentration (**Fig. 3C**). Re-expression of A3B in *A3B*-null U2OS cells partially restored pan-nuclear γH2AX accumulation. A similar pattern was observed following Prexasertib treatment, with A3B-dependent differences most evident at the IC_50_ concentration and attenuated at the IC_95_ concentration (**Fig. 3D**). Together, these results indicated that A3B enhances CHK1 inhibitor-associated pan-nuclear γH2AX accumulation, with a stronger effect in JHOC5 cells than in U2OS cells.

### Endogenous A3B impacts the cell cycle response to CHK1 inhibition

CHK1 inhibition can impair cell-cycle progression by promoting replication stress and accumulation of catastrophic DNA damage (47). To further investigate the consequences of CHK1 inhibition in cells with high endogenous A3B, JHOC5 and U2OS cells were stained with PI and analyzed by DNA content profiling after 24 h treatment with GDC-0575 or Prexasertib. In vehicle treated JHOC5 cells, A3B-proficient and *A3B*-null cells showed similar cell-cycle distributions (**Fig. 4A**). However, following treatment with GDC-0575 or Prexasertib, A3B-expressing JHOC5 cells exhibited dose-dependent alterations in DNA-content distribution, leading to cells accumulating in S-phase as expected. In contrast, *A3B*-null cells retained profiles more similar to vehicle-treated controls, with clear distinctions between G1, S, and G2/M phase, with increased accumulation of cells in G2/M phase. Re-expression of A3B in *A3B*-null cells restored the drug-induced redistribution pattern like that observed in A3B-proficient JHOC5 cells (**Fig. 4A**).

**Figure 4.**
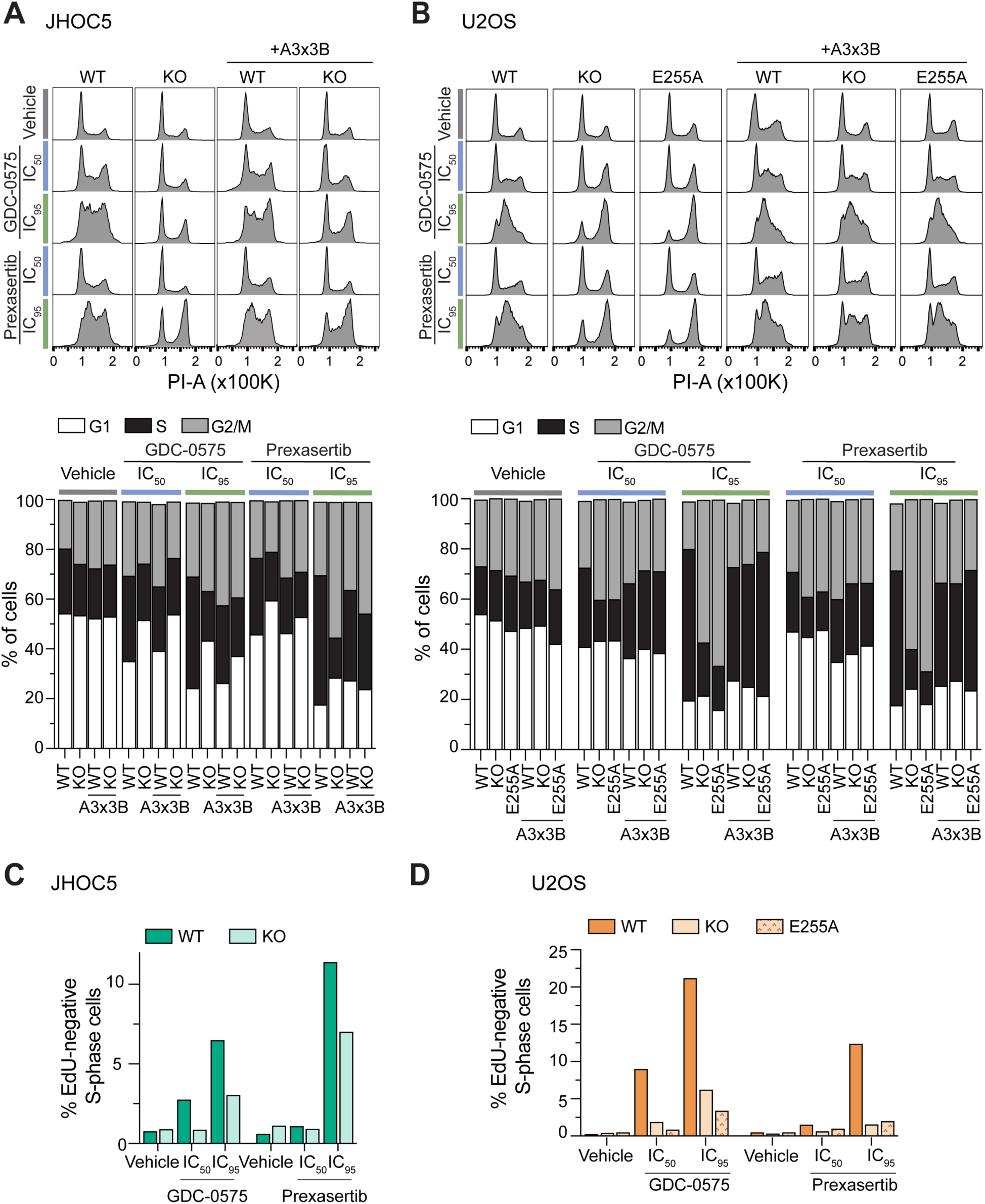
Endogenous A3B modulates cell cycle response to CHK1 inhibitors. **A**, Representative DNA content histograms (PI staining intensity) of JHOC5 *A3B* WT and *A3B* KO cells, with or without ectopic A3×3B expression, following treatment with vehicle, GDC-0575 at IC_50_ (0.15 µM) or IC_95_ (0.25 µM) concentrations, or Prexasertib at IC_50_ (0.005 µM) or IC_95_ (0.025 µM) concentrations. The histogram below shows a quantification of the cell cycle stages (G1, S, G2/M) for each of the indicated conditions. **B**, Representative DNA content histograms (PI staining intensity) of U2OS *A3B* WT and *A3B* KO cells, with or without ectopic A3×3B expression, following treatment with vehicle, GDC-0575 at IC_50_ (0.085 µM) or IC_95_ (0.25 µM) concentrations, or Prexasertib at IC_50_ (0.003 µM) or IC_95_ (0.0065 µM) concentrations. The histogram below shows a quantification of the cell cycle stages (G1, S, G2/M) for each of the indicated conditions. **C** and **D**, Quantification of the percentage of EdU-negative staining S-phase cells for JHOC5 and U2OS, respectively, following treatment with vehicle control, GDC-0575, or Prexasertib at IC_50_ or IC_95_ concentrations.

Similar experiments were done in U2OS cells, including A3B WT, A3B KO, and catalytically inactive A3B E255A knock-in cells. As in A3B WT JHOC5 cells, CHK1 inhibition with either GDC-0575 or Prexasertib led to dose-dependent changes in DNA content profiles (**Fig. 4B**). Again, A3B-proficient cells preferentially accumulated in S-phase after CHK1 inhibition. In contrast, *A3B*-null and A3B-E255A cells showed a reduced shift toward this aberrant S-phase profile, indicating that this response depends, at least in part, on A3B catalytic activity. The re-expression of A3B restored the CHK1 inhibitor-induced redistribution pattern in *A3B*-null cells and increased the response in E255A cells, further supporting a functional relationship between A3B activity and altered cell-cycle progression following CHK1 inhibition.

Since PI staining alone cannot distinguish between actively replicating cells versus cells with stalled replication, EdU incorporation in combination with PI was used to quantify EdU-negative S-phase cells. These cells have partially replicated DNA, but are not currently synthesizing new DNA, since most replication forks have stalled or arrested (53,54). In JHOC5 cells, CHK1 inhibition increased the fraction of EdU-negative S-phase cells more strongly in A3B-proficient than in *A3B*-null cells, with the largest increase observed after Prexasertib treatment at IC_95_ (**Fig. 4C**). A similar pattern was observed in U2OS cells, where GDC-0575 and Prexasertib increased EdU-negative S-phase cells A3B WT cells, whereas A3B KO and A3B E255A cells showed reduced accumulation of this population (**Fig. 4D**). As a result, these data indicated that, after CHK1 inhibition, A3B-proficient cells contained a larger EdU-negative S-phase population than *A3B*-null or E255A cells. This population has S-phase DNA content but no detectable EdU incorporation, consistent with impaired DNA synthesis.

## Discussion

In this study, we show that endogenous A3B confers a selective vulnerability to CHK1 inhibition. Using two cancer cell lines with pathologically high endogenous A3B expression and their *A3B*-null counterparts, we demonstrate that A3B-expressing cells are consistently more sensitive to two chemically distinct CHK1 inhibitors, GDC-0575 and Prexasertib. This sensitivity is apparent after short-term treatment, and it becomes even more pronounced in colony formation assays, indicating that sustained CHK1 inhibition is detrimental to cancer cell viability when A3B expression is high. The effect requires A3B catalytic activity, as a catalytically inactive mutant phenocopies genetic ablation, whereas re-expression of A3B in *A3B*-null cells restores sensitivity. Mechanistically, CHK1 inhibition drives the accumulation of pan-nuclear γH2AX in A3B-high cancer cells and causes a marked disruption of the cell cycle, whereas *A3B*-null cells retain a more clearly defined cell cycle profile. Consistent with this, upon CHK1 inhibition of high A3B-expressing cells, a much smaller percentage of S-phase cells stain positive for EdU, indicative of stalled DNA replication forks. These findings strongly indicate that A3B-mediated single-stranded DNA cytosine deamination increases replication stress and, upon CHK1 inhibition, triggers cell death.

It is interesting that the clear cell ovarian cancer cell line JHOC5 exhibits a much stronger A3B-dependent sensitivity to CHK1 inhibition than the U2OS cell line. U2OS is derived from an osteosarcoma (55), and this cell line is notable for maintaining its telomeres through the alternative lengthening of telomeres (ALT) pathway which is a telomerase-independent mechanism driven by homology-directed, break-induced DNA synthesis at telomeres (56). Because ALT-positive cells rely on elevated homologous recombination and specialized replication-fork repair this could potentially influence U2OS cells response A3B-mediated DNA damage. An increased capacity for homologous recombination might provide a buffer against A3B-mediated DNA deamination, by offering an efficient mechanism to repair A3B-catalyzed single- and double-stranded DNA breaks. This compelling mechanistic relationship is most recently evidenced by a profound A3B sensitivity in BRCA2-deficient cells (46). Thus, cancer cell lines and tumors that have normal (or downregulated) levels of homologous recombination and upregulated levels of endogenous A3B, such as JHOC5, might be especially susceptible to CHK1 inhibition.

Another notable finding from our study is that, despite the close functional relationship between ATR, CHK1, and WEE1, inhibition of ATR or WEE1 produces weaker effects in comparison to direct CHK1 inhibition. ATR inhibition reduces colony formation in the ovarian cancer cell line JHOC5 and in osteosarcoma U2OS, but the effect is not dependent on A3B expression. In comparison, WEE1 inhibition produces modest A3B-dependent differences in both lines. Prior work has emphasized ATR as a principal vulnerability created by APOBEC3 activity where ectopic expression of A3A and A3B sensitizes cells to ATR inhibition (34), and high A3B has been reported to confer sensitivity to both ATR and CHK1 inhibitors in an R-loop–dependent manner (33). Ectopic overexpression or doxycycline-inducible A3B systems can produce expression levels and temporal patterns that differ from those of endogenous A3B. Expression of the endogenous *A3B* gene is cell cycle regulated and expressed preferentially during the G2/M phase and repressed in G0/G1 (44,57). Constitutive expression therefore places A3B protein and activity in cell cycle phases where it is normally absent, which may drive more uracil lesions, abasic sites, DNA breaks, and replication stress, and in turn a potentially amplified ATR-directed checkpoint dependency. Consistent with this possibility, dox-inducible A3B expression alone increases the fraction of EdU-negative S-phase cells in HEK-293 cells, which lack detectable endogenous A3B (32). In our isogenic models, by contrast, A3B-proficient and *A3B*-null cells are indistinguishable in EdU-negative content at baseline, and differences only emerge upon inhibition of CHK1. These findings suggest that the levels of A3B in several human cancer types and cell lines may not produce sufficient replication stress to affect cell survival, but these levels have the potential to become toxic upon CHK1 inhibition.

## Authors’ contributions

**B. Stefanovska:** Conceptualization, formal analysis, investigation, methodology, project administration, funding acquisition, visualization, writing - original draft. **B.C. Troness:** Formal analysis, investigation, methodology and visualization. **C.D. Mullally:** Investigation and methodology. **B. de la Peña Avalos:** Investigation. **M.A. Ibrahim:** Investigation. **Y. Chen:** Investigation. **E. Fanunza:** Investigation. **M.A. Carpenter:** Investigation and methodology. **R.S. Harris:** Conceptualization, investigation, methodology, project administration, supervision, funding acquisition, writing - original draft. **All authors:** writing – review & editing.

## Acknowledgements

We thank Alan Ashworth, Sofia Henderson, Yu-Hsiu Tony Lin, and members of the Harris laboratory for constructive feedback, and Emma Maudal for early experiments with DNA repair inhibitors. This work was supported by NCI P01-CA234228, NCI P50-CA247749, and a Recruitment of Established Investigators Award from the Cancer Prevention and Research Institute of Texas (CPRIT RR220053). BS was supported in part by a grant from the Ovarian Cancer Research Alliance (OCRA Mentored Investigator Grant 812337). CDM and YTL received salary support from the South Texas Medical Scientist Training Program (NIGMS T32-GM113896 and T32-GM145432) and the Epigenetics, DNA Repair, and Genomics (EDGe) Training Program (NCI T32-CA279363). We thank the technical support from the Mays Cancer Center Drug Discovery Shared Resource (consisting of the Center for Innovative Drug Discovery and Target Discovery Core) at University of Texas Health San Antonio, which is supported by CPRIT Core Facility Award RP250601 and National Cancer Institute Cancer Center support grant P30 CA054174. RSH is an investigator of the Howard Hughes Medical Institute, a CPRIT scholar, and the Ewing Halsell President’s Council Distinguished Chair at the University of Texas Health San Antonio. Generative AI was used solely to correct grammar and improve language clarity, and any suggested changes were reviewed manually by the authors. The authors have no competing interests to declare.

