## Supplemental Table 1 and Figs 1-3 for "Endogenous APOBEC3B Promotes CHK1 Inhibitor Sensitivity"

Running Title: APOBEC3B and CHK1 interaction

[*]Equal Contributions

Supplement information: Supplementary Table 1; Supplementary Figs. 1-3

**Supplementary Table 1.**

| **Antibodies** | | |
|---|---|---|
| Protein target | Manufacturer | Catalog number |
| Chk1 | Cell Signaling Technology | 2360S |
| p-Chk1 S345 | Cell Signaling Technology | 2348S |
| p-Chk1 S296 | Cell Signaling Technology | 2349S |
| γH2AX S139 | Millipore Sigma | 05-636 |
| A3A/B/G | In-house production | 5210-87-13 |
| α-tubulin | Millipore Sigma | T5168 |
| Goat polyclonal secondary anti-rabbit IgG, HRP-linked | Cell Signaling Technology | 7074 |
| Goat anti-mouse IgG IRDye 680LT | LI-COR | 926-68020 |
| **Primers** | | |
| Description | Oligo sequence (5'-to-3') | |
| Outer forward primer for *A3B* KO | GCCCTTCCAGATAGAGGGCAAGAGACAG | |
| Outer reverse primer for *A3B* KO | CTGAGATGAAGAAGCGGGAAGCAGTCAGG | |
| Inner forward primer for *A3B* KO | TCCTGCTCCCCCTCTCAGAGCATC | |
| Inner reverse primer for *A3B* KO | GGCTGTCAGTTGCAGTCAGTGCCAG | |
| Outer forward primer for A3B E255A | TGCATCCCCTCTGATGGAA | |
| Outer reverse primer for A3B E255A | CCCAGTTCTCTTTCTTTCGGAAAT | |
| Inner forward primer for A3B E255A | CTAATAACCGGGGTTTTGG | |

| | |
|---|---|
| Inner reverse primer for A3B E255A | CCCGCAGCATTTGCAGCGC |
| **gRNA sequences (5’-to-3’)** | |
| A3B KO | ATAGTCCATGATCTTCACGC |
| A3B E255A | AGAAGCGCAGCTCCGCATGG |
| A3B E255A ssODN | CTAAGAATCTTCTCTGTGGCTTTTACGGCCGCCATGCGG AGCTGCGCTTCTTGGACCTGGTTCCTTCTTTGCAGTTGG |

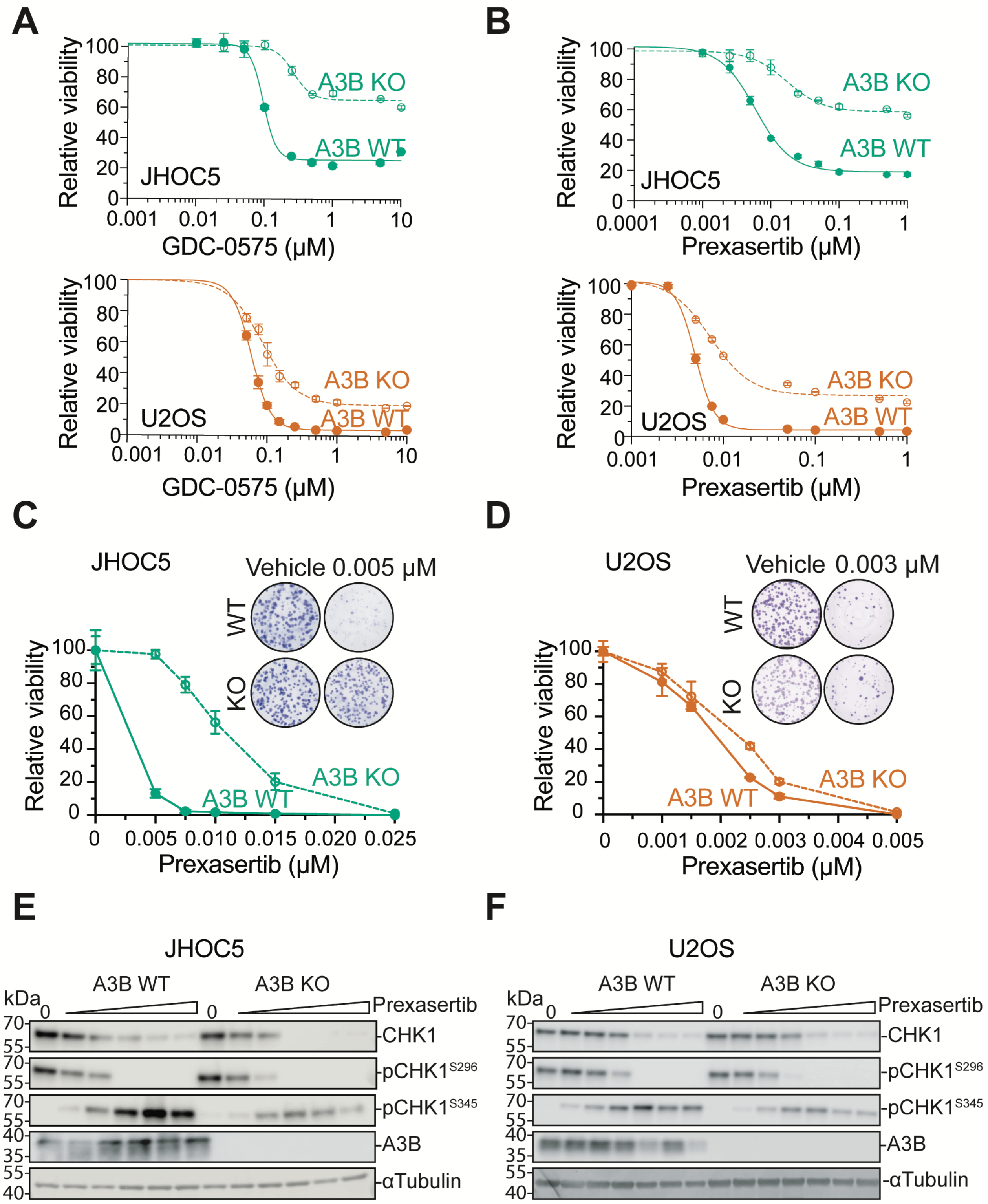
A
Relative viability
GDC-0575 (μM)
JHOC5
U2OS
A3B KO
A3B WT
B
Prexasertib (μM)
C
JHOC5
Vehicle 0.005 μM
WT
KO
D
U2OS
Vehicle 0.003 μM
E
JHOC5
A3B WT
A3B KO
Prexasertib
kDa
CHK1
pCHK1S296
pCHK1S345
A3B
αTubulin
F
U2OS

**Supplementary Figure 1.** APOBEC3B-proficient cells are hypersensitive to the CHK1 inhibitor Prexasertib.

**A**, Dose-response analysis of JHOC5 and U2OS *A3B* WT and *A3B* KO cells treated with the CHK1 inhibitor GDC-0575. Relative viability was measured with CellTiter-Glo after drug treatment and normalized to vehicle-treated controls. Data are mean ± SD, n = 3 independent experiments. (P-value <0.001 for WT vs KO by extra sum-of-squares F-test)

**B**, Dose-response analysis of JHOC5 and U2OS *A3B* WT and *A3B* KO cells treated with the CHK1 inhibitor Prexasertib. Relative viability was measured with CellTiter-Glo after drug treatment and normalized to vehicle-treated controls. Data are mean ± SD, n = 3 independent experiments. (P-value <0.01 for WT vs KO in JHOC5 and P-value <0.001 in U2OS by extra sum-of-squares F test)

**C** and **D**, Dose-response analysis of *A3B* WT versus KO JHOC5 and U2OS cells, respectively, following treatment with the indicated concentrations of the CHK1 inhibitor Prexasertib. Relative viability was measured after drug treatment and normalized to vehicle-treated controls. Representative colony-formation images are shown. Data are mean ± SD, n = 3 independent experiments. (P-value <0.0001 for WT vs KO in JHOC5 and P-value =0.04 in U2OS by extra sum-of-squares F-test).

**E**, Immunoblot analyses of CHK1 signaling in *A3B* WT and KO JHOC5 following treatment with 0, 0.00005, 0.0005, 0.005, 0.05, and 0.5 μM Prexasertib.

**F**, Immunoblot analyses of CHK1 signaling in *A3B* WT and KO U2OS following treatment with 0, 0.00001, 0.0001, 0.001, 0.01, 0.1, and 1 μM Prexasertib.

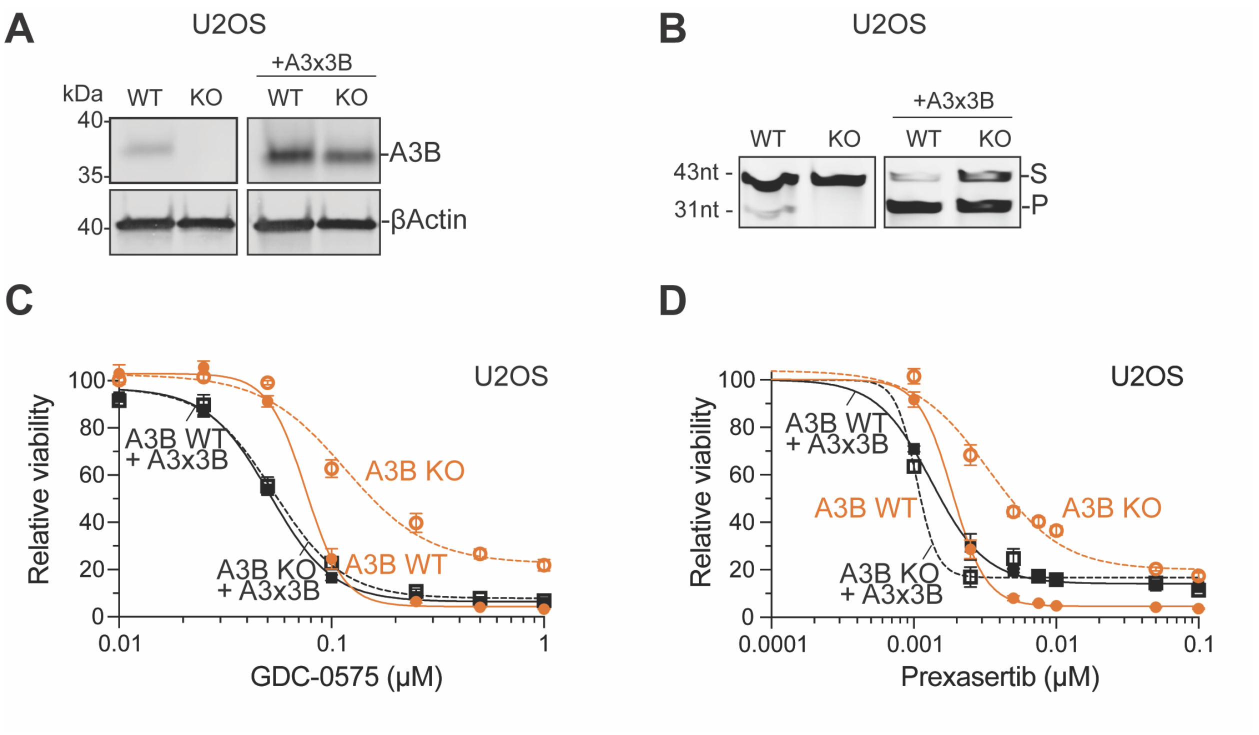


**Supplementary Figure 2.** APOBEC3B catalytic activity promotes sensitivity to CHK1 inhibition.

**A**, Anti-A3B immunoblot demonstrating CRV-mediated expression of exogenous A3B in WT and KO U2OS cells

**B**, C-to-U activity assay demonstrating CRV-mediated expression of exogenous A3B in WT and KO U2OS cells (S, substrate; P, product).

**C** and **D**, Dose-response analysis of untransduced or CRV-transduced *A3B* WT versus *A3B* KO U2OS cells following treatment with GDC-0575 and Prexasertib, respectively. Relative viability was quantified using CellTiter-Glo 72 h after drug treatment (normalized to vehicle-treated controls; mean ± SD of 3 independent experiments; P-value <0.0001 for WT vs KO by extra sum-of-squares F-test; all other comparisons are not significant).

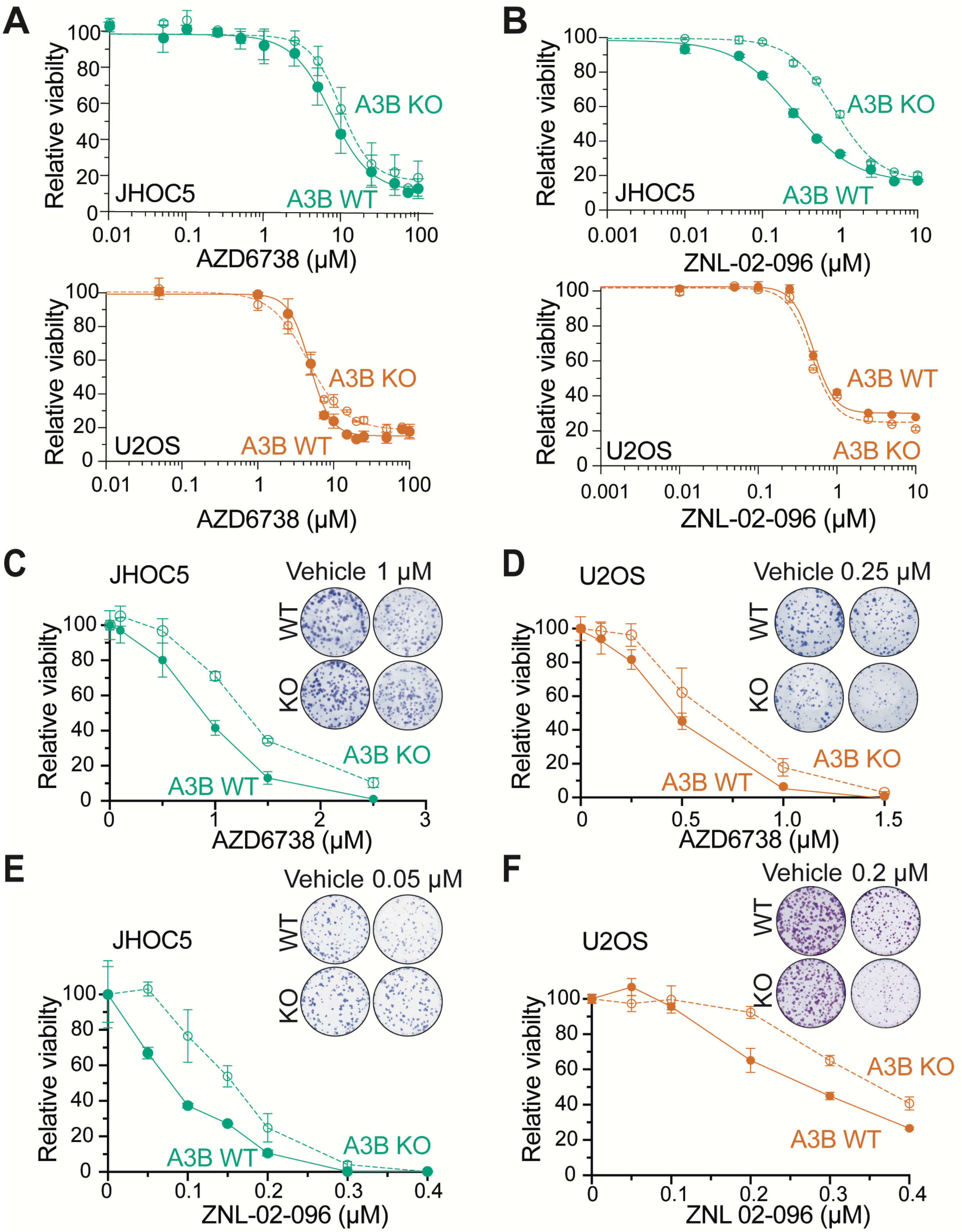
A
Relative viabilty
JHOC5
A3B KO
A3B WT
AZD6738 (μM)
U2OS
B
ZNL-02-096 (μM)
C
Vehicle 1 μM
WT
KO
D
Vehicle 0.25 μM
E
Vehicle 0.05 μM
F
Vehicle 0.2 μM
ZNL 02-096 (μM)

**Supplementary Figure 3.** A3B-dependent responses to ATR and WEE1 targeting are weaker and context dependent

**A**, Dose-response analysis of JHOC5 and U2OS *A3B* WT and *A3B* KO cells treated with the ATR inhibitor AZD6738. Relative viability was measured with CellTiter-Glo after drug treatment and normalized to vehicle-treated controls. Data are mean ± SD, n = 3 independent experiments. (WT vs KO comparisons are non-significant by extra sum-of-squares F-test)

**B**, Dose-response analysis of JHOC5 and U2OS *A3B* WT and *A3B* KO cells treated with the WEE1 degrader ZNL-02-096. Relative viability was measured with CellTiter-Glo after drug treatment and normalized to vehicle-treated controls. Data are mean ± SD, n = 3 independent experiments. (P-value <0.001 for WT vs KO in JHOC5 and non-significant in U2OS by extra sum-of-squares F test)

**C** and **D**, Dose-response analysis of *A3B* WT versus KO JHOC5 and U2OS cells, respectively, following treatment with the indicated concentrations of the ATR inhibitor AZD6738. Relative viability was measured after drug treatment and normalized to vehicle-treated controls. Representative colony-formation images are shown. Data are mean ± SD, n = 3 independent experiments. (P-value <0.01 for WT vs KO by extra sum-of-squares F-test).

**E** and **F**, Dose-response analysis of *A3B* WT versus KO JHOC5 and U2OS cells, respectively, following treatment with the indicated concentrations of the WEE1 degrader ZNL-02-096. Relative viability was measured after drug treatment and normalized to vehicle-treated controls. Representative colony-formation images are shown. Data are mean ± SD, n = 3 independent experiments. (P-value <0.0001 for WT vs KO in JHOC5 and non-significant in U2OS by extra sum-of-squares F-test).
